# A programmable arthritis-specific receptor for guided articular cartilage regenerative medicine

**DOI:** 10.1101/2024.01.31.578281

**Authors:** Bonnie L. Walton, Rebecca Shattuck-Brandt, Catherine A. Hamann, Victoria W. Tung, Juan M. Colazo, David D. Brand, Karen A. Hasty, Craig L. Duvall, Jonathan M. Brunger

## Abstract

**Objective:** Investigational cell therapies have been developed as disease-modifying agents for the treatment of osteoarthritis (OA), including those that inducibly respond to inflammatory factors driving OA progression. However, dysregulated inflammatory cascades do not specifically signify the presence of OA. Here, we deploy a synthetic receptor platform that regulates cell behaviors in an arthritis-specific fashion to confine transgene expression to sites characterized by cartilage degeneration.

**Methods:** An scFv specific for type II collagen (CII) was used to produce a synthetic Notch (synNotch) receptor that enables “CII-synNotch” mesenchymal stromal cells (MSCs) to recognize CII fibers exposed in damaged cartilage. Engineered cell activation by both CII-treated culture surfaces and on primary tissue samples was measured via inducible reporter transgene expression. TGFβ3-expressing cells were assessed for cartilage anabolic gene expression via qRT-PCR. In a co-culture with CII-synNotch MSCs engineered to express IL-1Ra, ATDC5 chondrocytes were stimulated with IL-1α, and inflammatory responses of ATDC5s were profiled via qRT-PCR and an NF-kB reporter assay.

**Results:** CII-synNotch MSCs are highly responsive to CII, displaying activation ranges over 40-fold in response to physiologic CII inputs. CII-synNotch cells exhibit the capacity to distinguish between healthy and damaged cartilage tissue and constrain transgene expression to regions of exposed CII fibers. Receptor-regulated TGFβ3 expression resulted in upregulation of *Aca*n and *Col2a1* in MSCs, and inducible IL-1Ra expression by engineered CII-synNotch MSCs reduced pro-inflammatory gene expression in chondrocytes.

**Conclusion:** This work demonstrates proof-of-concept that the synNotch platform guides MSCs for spatially regulated, disease-dependent delivery of OA-relevant biologic drugs.

## INTRODUCTION

Osteoarthritis (OA) is a chronic, progressively degenerative joint disease for which there are no clinically approved disease-modifying osteoarthritis drugs (DMOADs). While OA has been classically considered a “wear and tear” disease, it is now understood that mechanical, cellular, and soluble factors cumulatively contribute to its progression. The hallmark of disease progression remains the catabolism of articular cartilage, which is exacerbated by dysregulated inflammatory cascades, immune cell infiltration, and increased activity of extracellular matrix (ECM)-degrading proteases, such as matrix metalloproteinases (MMPs) and aggrecanases (1–4). These features result in overall loss of cartilage volume, subchondral bone remodeling, synovial thickening, and lymphatic dysregulation. Noninvasive OA treatments are palliative in nature, with an emphasis on pain management and slowed disease progression. Later stage treatments resort to surgical intervention such as autologous chondrocyte implantation (5, 6), microfracture (7), or total joint replacement (8). Many of these surgical procedures aim to introduce progenitor cells to the arthritic lesion for neotissue development, as native joint-resident cells do not adequately regenerate articular cartilage with the same biomechanical properties(9). Prospective OA cell therapies, such as autologous chondrocyte or mesenchymal stromal cell (MSC) transplantation, may offer advantages because cells can integrate multiple inputs to perform a variety of outputs ranging from mitigating inflammatory cascades (10–12) to stimulating cartilage regeneration (13, 14). As such, compared to traditional biologic drugs, cells serve as more versatile agents to coordinate repair of damaged tissue. However, while these approaches offer some clinical benefit, outcomes decline over time, primarily due to cell death in the degenerative environment, as well as the development of fibrocartilage (15–18). In addition, most traditional cell-based therapies are not capable of selectively responding to OA pathology for directed regenerative functions.

Synthetic biology, the field that attempts to develop new biological systems with user-defined input/output relationships, offers an additional layer of control over the behavior of engineered cells. Synthetic biology techniques to create “designer” cell therapies have facilitated the development of several medical breakthroughs such as chimeric antigen receptor (CAR) T cell cancer immunotherapy (19, 20). The CAR directs T cell activity toward cells displaying antigens of choice, typically tumor-related antigens, by programming a mimetic of the T cell receptor with a recognition motif specific for such antigens. In a similar way, an ideal cell engineered for the treatment of OA would be programmed to survey the joint microenvironment and activate localized, feedback-regulated drug production to mitigate joint-resident inflammation and promote cartilage repair only in the context of disease. However, genetic engineering strategies to overcome the limitations of cell therapies for OA have primarily focused on the constitutive or drug-inducible expression of therapeutic factors to mitigate the arthritic environment (11, 21–23). These approaches override native microenvironmental inputs and offer no localized control over the expression of disease-modifying factors. Factors such as mechanical degradation, post-traumatic injury, and genetic predisposition ultimately lead to the upregulation of pro-arthritic biomarkers that may serve as OA-specific cell engineering targets. Recent advances in synthetically regulated cells for OA aim to rewire native signaling pathways to respond to these targets for autoregulated expression of biologic drugs (24–26). However, these approaches have resulted in cells that respond to pleiotropic features not exclusive to arthritis progression. Further, these artificial signaling pathways do not instruct cells to implement regenerative behaviors localized to the arthritic joint. As such, these therapies are not optimized for feedback-controlled expression of therapeutic factors specific to OA pathology.

We present a strategy to engineer cells to detect OA pathology via the synthetic Notch (synNotch) receptor. SynNotch is based on the native juxtacrine Notch receptor, which activates downstream gene expression upon binding to an immobilized ligand (27, 28). Like Notch, synNotch activation depends on a 4-12 pN tensile force generated via mechanical cell-ligand interactions within a niche (29, 30). This dependence on mechanical force qualifies synNotch as a useful tool for regulating highly localized gene expression programs in response to receptor inputs (27, 31, 32). By exchanging Notch’s extracellular domain with single chain antibodies and intracellular domain with a synthetic transcription factor, synNotch receptors that produce user-defined sense/response behaviors can be created. We previously characterized a monoclonal antibody against type II collagen (CII), known as mAbCII, that specifically recognizes epitopes of CII unmasked during cartilage matrix degradation and identifies early OA lesions (33–35). MAbCII has been used as a theranostic moiety to selectively identify damaged cartilage tissue and as a homing device to deliver short-interfering RNA (siRNA) cargo to arthritic joints (35, 36). Here, we speculated that we could engineer cells to directly probe for cartilage matrix degradation and implement a pre-selected, pro-regenerative response via the synNotch signaling channel. Thus, we programmed the recognition motif of synNotch with a single chain variable fragment (scFv) derived from the light and heavy variable domains from the mAbCII antibody. Using reporter transgenes, we characterized the specificity, sensitivity, and dynamic range of this new synNotch receptor, CII-synNotch, that incorporates the mAbCII recognition domain. Gene circuits were then constructed to regulate pro-anabolic and anti-inflammatory factors that enable regenerative functions of CII-synNotch cells. Results presented here highlight the capacity to engineer synNotch cells that directly probe for cartilage matrix degradation and then implement defined, feedback-regulated responses based on this local, pathology-dependent assessment of cartilage status.

## METHODS

### SynNotch receptor and “payload” transgene vectors

To construct the CII-synNotch receptor, a gene fragment encoding the CD8α signal peptide, c-myc epitope tag and the variable heavy chain from mAbCII linked to the variable light mAbCII chain was procured from GeneArt (ThermoFisher). This fragment was cloned into a lentiviral expression plasmid we have previously described (31) encoding the Notch1 core linked to the tTA transcription factor using NEBuilder HiFi DNA Assembly (New England Biolabs). The resultant plasmid encoded constitutive expression of CII-synNotch receptor from an EF1α promoter and production of the puromycin N-acetyl-transferase transgene via the constitutive PGK promoter for antibiotic selection. As a control receptor, we used the anti-GFP LaG16 nanobody-based synNotch receptor we have previously reported (31). We refer to the downstream gene circuit activated by synNotch as the “payload’ expression cassette. The luciferase+mCherry and SEAP payload constructs were previously described (31) and contained transgenes downstream of the TRE3G tet-responsive promoter as well as constitutive, PGK-driven tagBFP or iRFP720, as indicated. TGFβ3 and IL-1Ra coding sequences were similarly cloned downstream of TRE3G tet-responsive promoter via NEBuilder HiFi DNA Assembly in vectors encoding PGK-driven tagBFP.

### Lentivirus production

Second-generation lentivirus was produced by co-transfecting the pCMV-dR8.91 gag/pol vector (1.5 μg), the pMD2.G envelope vector (0.6 μg, a gift from Didier Trono, Addgene #12259) and the receptor- or payload-encoding viral expression plasmids (2.0 μg) into Lx293T cells (Clontech) in 6-well plates with Lipofectamine 3000 (Thermo Fisher L3000001). To produce NF-κB reporter vector, a viral expression plasmid encoding mKate2 downstream of tandem NF-κB binding sites (37) (Addgene #105173, a gift from Timothy Lu) was co-transfected with gag/pol and env vectors. Virus production medium contained DMEM supplemented with L-glutamine and sodium pyruvate and 10% heat-inactivated FBS (Gibco). Viral supernatants collected roughly 48- and 72-hours post-transfection were sterile-filtered through a 0.45-micron filter and concentrated in 100kDa molecular weight cutoff filters.

### Murine mesenchymal stromal cells

Bone marrow-derived mesenchymal stromal cells from C57BL/6 mice (Cyagen) were cultured in MEMα supplemented with GlutaMAX (Gibco) in 15% fetal bovine serum (Gibco). Engineered mMSCs were generated via forward transduction by resuspending the concentrated receptor and payload vectors at a 2:1 volumetric ratio in 2ml of mMSC sub-cultivation medium containing 4 μg/ml polybrene, which was added to the cell monolayer. Viral medium was removed and replaced with sub-cultivation medium after 16 hours. Prior to use, cells were sorted at the Vanderbilt University Medical Center Flow Cytometry Shared Resource Core. Cells were immunolabeled using an Alexa Fluor 647 anti-c-myc antibody (Cell Signaling, 9B11), which binds the c-myc epitope tag on the N-terminus of the receptor to identify synNotch receptor^+^ cells.

### ATDC5 chondrocytes

Immortalized mouse ATDC5 chondrocytes (Sigma-Aldrich) were cultured in DMEM/F12 containing L-glutamine (Gibco) supplemented with 10% FBS. NF-κB reporter ATDC5 cells were produced by supplementing ATDC5 maintenance medium with viral supernatant, which was concentrated and re-suspended in ATDC5 sub-cultivation medium, at a 1:1 ratio and 4 μg/ml polybrene for overnight transduction.

### Preparation of collagen decorated surfaces

Type I and type II collagen stocks were extracted and purified from bovine skin and articular cartilage, respectively using a differential salt precipitation-based methodology. Collagen was solubilized in Jerry Gross buffer (127 mM K_2_HPO4, 3.4 mM KH_2_PO_4_, pH 7.6) at 4°C for at least 24 hours before use. After solubilization, collagen was stored at −80°C until further use. For all synNotch characterization studies, collagen was thawed and diluted to indicated concentrations to coat the bottom of tissue culture treated plates. After adding collagen, plates were sealed with parafilm and placed at 4°C for approximately 16 hr or overnight. Excess, unbound collagen was aspirated from wells prior to plating cells for experiments.

### Cartilage tissue sample preparation

Whole tissue samples were obtained from the femoral cartilage of adolescent male Yorkshire pigs, approximately 1 hr after being sacrificed for unrelated studies. Briefly, 3 mm biopsy punches (Integra) were used to puncture the cartilage and create the plugs used for whole sample experiments. Plugs were washed for two hours in 2x antibiotic-antimycotic (Gibco) in DPBS before long-term storage at −80°C. When using tissue cryosections, cartilage plugs were embedded in optimal cutting temperature (OCT) compound and sectioned into 10 μm thick sections using a cryotome at the Vanderbilt University Translational Pathology Shared Resource core facility. Sections were stored at −80°C until use.

### Luciferase detection to quantify synNotch activity

For activation experiments on passively adsorbed collagen, synNotch-luciferase cells were seeded at a density of 39,000 cells/cm^2^ on the indicated surfaces treated with or without types I or II collagen. Detection of inducible luciferase was assessed after lysing cells via the addition of BrightGlo (Promega) 72 hours later, unless otherwise specified. Luminescent signal was measured on a Tecan Infinite M1000 Pro plate reader.

### SEAP detection to measure synNotch activity

A subset of cartilage tissue explants was incubated in TrypLE express for one hour at 37C before aspirating the TrypLE, washing explants in PBS containing 10% FBS and antibiotic/anti-mycotic. Control explants were incubated in PBS containing 10% FBS with antibiotic/anti-mycotic. SynNotch-SEAP MSCs were plated on explants (100,000 cells in 5 μl medium) and allowed to adhere for 1 hour prior to adding culture medium containing 1x antibiotic-antimycotic. SEAP production was analyzed by collecting medium and running a chemiluminescent assay on samples (Takara Bio) using a Tecan Infinite M1000 Pro plate reader.

### SynNotch activation on cartilage cryosections

Cryosections were brought to room temperature, washed in DPBS containing 2x antibiotic-antimycotic, and transferred to a well-plate also containing DPBS containing 2x antibiotic-antimycotic. After these washes, sections were incubated in noted concentrations of trypsin-EDTA (Gibco) for one hour of incubation at 37°C. Trypsin was then quenched in DPBS containing 5% FBS and 1x antibiotic-antimycotic and aspirated. Sections were washed in DPBS, and 10,000 cells were seeded in 50 µl directly to the surface of the tissue and incubated at 37°C for 45 minutes before 1x antibiotic-antimycotic-containing MSC medium was added to wells containing sections. Fluorescence microscopy experiments enabling identification of synNotch-driven mCherry expression were performed in tissue culture-treated plates. Samples prepared for luminescence assays were cultured on ultra-low attachment plates. To detect inducible luciferase expression, BrightGlo was added to the well for lysis of engineered cells, and the lysate was transferred to a new well prior to acquiring a luminescence reading. Cell metabolic activity was measured via CellTiter-Fluor (Promega) as a surrogate for cell number to account for potential differences in cell attachment to tissue vs. untreated or CII-decorated surfaces. Raw luminescence values were divided by CellTiter-Fluor values to calculate normalized luminescence.

### Trans-well MSC:ATDC5 co-culture experiment

MSCs were plated on surfaces prepared with 25 μg/ml adsorbed CII, as previously described, in the bottom of a 24-well tissue culture treated plate in 0.5 ml MSC medium. ATDC5s were plated in an upper chamber on a 3-micron porous insert in 0.2 ml ATDC5 medium. After three days of synNotch activation, medium from the upper chamber was removed and replaced with ATDC5 medium containing 0 or 0.1 ng/ml IL-1α (STEMCELL Technologies). Following two additional days of incubation, inserts containing ATDC5s were transferred to fresh wells, where ATDC5 cells were lysed for mRNA isolation and subsequent gene expression analysis.

### Flow cytometry

Cells were washed with 1x DPBS and dissociated with TrypLE Express (Gibco) before being quenched with DPBS containing 5% FBS. To track fluorescent expression, samples were passed through a CellStream analytical flow cytometer. Analysis was performed in FlowJo and all data shown is gated on the basis of singlet, live cell events, and plotted as median fluorescence intensity of the respective fluorescence marker in gated populations.

### ELISA

In all ELISA experiments, CII-synNotch cells were activated on cell culture surfaces treated with 25 μg/ml CII or plated on untreated surfaces. Medium was collected from samples after 72 hr of culture. DuoSet Human IL-1Ra or TGFβ3 ELISA detection kits (R&D) were used with the DuoSet Ancillary Reagents kit for detection of secreted protein (R&D). For the TGFβ3 ELISA, latent TGFβ3 in the medium was activated for detection prior to data acquisition.

### Gene expression analysis

Isolation of mRNA was achieved after lysing cells using PureLink Mini Kits (Invitrogen). Reverse transcription for cDNA was achieved with SuperScript IV VILO Master Mix (Invitrogen). Normalizing the mass of cDNA across samples, quantitative PCR was performed with SYBR Green Master Mix. Primer pairs used for detection of gene expression are listed in Table 1. Relative gene expression was calculated with the delta-delta Ct method. Unless otherwise noted, all samples are compared to the *r18s* housekeeping gene and normalized to cells in no-treatment conditions.

**Table 1:**
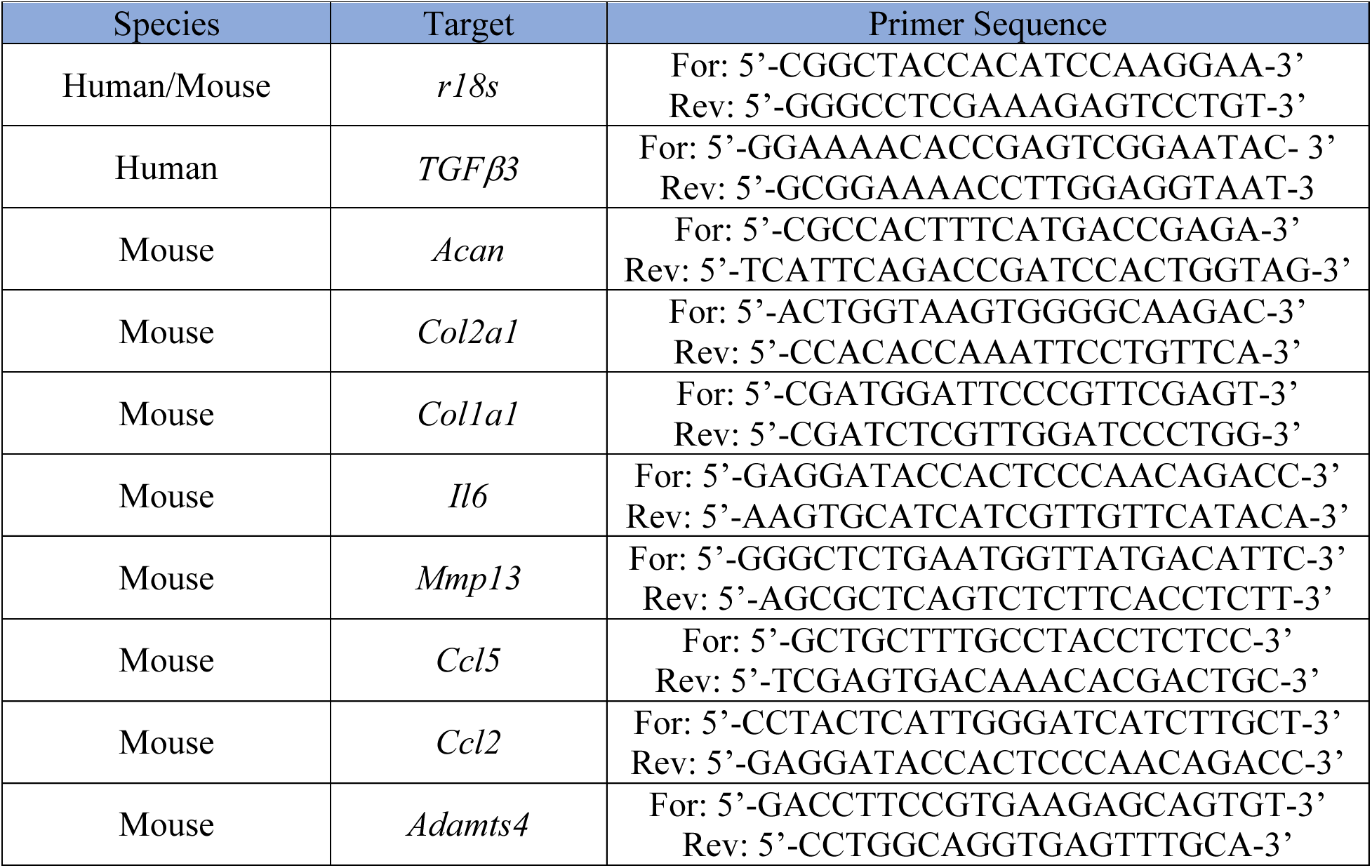
Sequences of oligonucleotides used as primers for qRT-PCR gene expression profiling.

### Fluorescence microscopy

All images were taken with a Leica Dmi8 epifluorescence microscope.

### Statistical analysis

All statistical analyses were performed with GraphPad Prism 10 software. Unless otherwise noted, plotted values represent group means ± standard error of the mean. With two comparisons, a Student’s t-test was performed with α=0.05. For multi-factorial comparisons, one- or two-way ANOVA with Tukey’s post-hoc analysis was performed, also with α=0.05.

## RESULTS

### Characterization of a type II collagen-specific synNotch receptor

We engineered a synNotch receptor with an extracellular domain composed of the light and heavy chains of the mAbCII antibody. We retained the Notch1 core shared amongst previously reported synNotch receptors (27, 28). As in our and others prior work (27, 31, 32), the tetracycline-controlled transactivator (tTA) served as the receptor intracellular domain and transcription factor that activates expression of payload transgenes encoded downstream of the tetracycline response element (TRE) promoter. Upon recognition of CII exposed after cartilage matrix damage, transmembrane cleavage of the Notch core leads to subsequent tTA translocation to the nucleus and TRE-driven transgene expression (**Figure 1A**). In this way, our CII-sensitive artificial receptor can activate orthogonal gene expression patterns, i.e., transcription programs that do not crosstalk with native signaling pathways (27).

**Figure 1:**
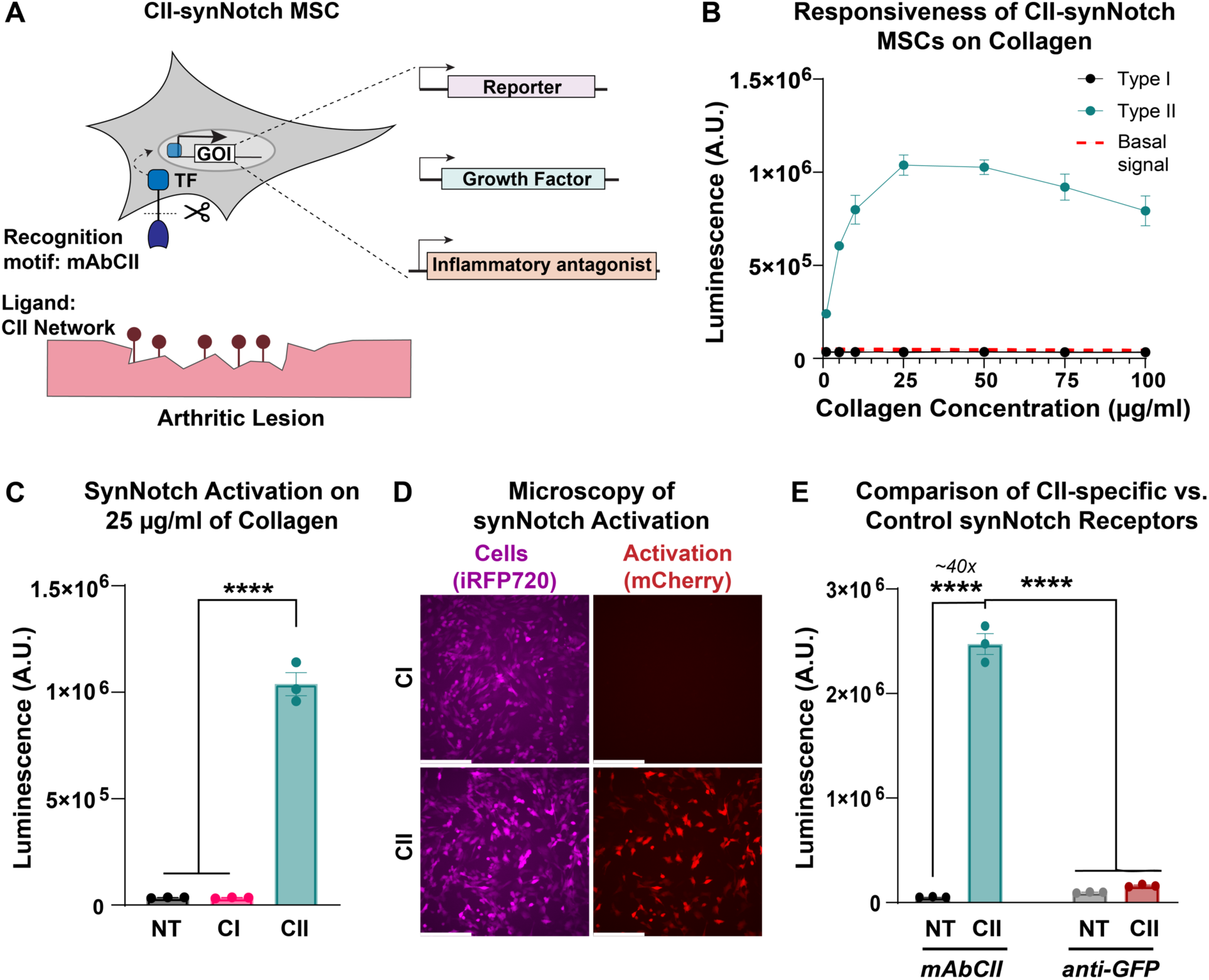
A mAbCII-programmed synNotch receptor licenses cells for potent and specific responses to type II collagen. **(A)** The type II collagen (CII)-sensitive antibody mAbCII serves as a synNotch recognition motif, rendering a new synNotch receptor, CII-synNotch. CII-synNotch facilitates cell recognition of CII fibers, leading to transmembrane cleavage of a synthetic transcription factor for versatile output responses. **(B)** CII-synNotch MSCs were plated on surfaces treated with solutions of varying concentrations of collagen I (CI) and CII (t=72hr). Red dashed line indicates basal signal from untreated cells. **(C)** Luminescence values of engineered cells on either 25 µg/ml CI or CII (t=72hr). **(D)** Fluorescence microscopy of constitutively iRFP^+^ CII-synNotch MSCs with inducible mCherry expression (t=72hr). **(E)** Activation of synNotch MSCs with either a mAbCII receptor or non-cognate (anti-GFP) receptor, measured via luciferase detection (t=72hr). Scale = 200 µm. Data: n=3 replicates, plotted as mean ± SEM. Statistical analysis: one-way ANOVA with Tukey’s post-hoc analysis, ****p < 0.0001. NT: No treatment.

Due to the widespread use of MSCs in arthritis therapy (11, 38–41), we opted to test the synNotch platform in primary MSCs. With a goal of applying these cells in future *in vivo* studies, we used murine bone marrow-derived MSCs. We first sought to characterize the specificity and sensitivity of the CII-synNotch receptor. Thus, we transduced MSCs with engineered lentiviral vectors encoding the CII-synNotch receptor as well as payload expression cassettes that enable regulated production of firefly luciferase and the mCherry fluorescent protein as reporter transgenes to monitor activation kinetics of the CII-synNotch signaling system. MSCs were plated on culture surfaces treated for passive adsorption with solutions of type I (CI) or type II (CII) collagen at varied concentrations (1-100 μg/ml). Surface treatment with CI resulted in similar levels of activation as cells cultured on control, not treated (NT) cell culture surfaces, as measured by luminescence activity (**Figure 1B-C**). However, surface treatment with CII yielded potent, dose-dependent activation of synNotch-driven transgene expression, which peaked in the 25 μg/ml surface treatment condition (**Figure 1B-C**). Fluorescence microscopy confirmed the potent activation levels quantified by the luminescence assay: while synNotch cells (labeled as iRFP^+^) are present in both CI and CII conditions, only cells on CII-treated surfaces demonstrate robust mCherry expression (**Figure 1D**). To demonstrate that CII-sensitivity was conferred by the mAbCII variable domains rather than a non-specific feature of synNotch, CII recognition was compared between CII-synNotch MSCs and MSCs engineered with a GFP-sensitive, LaG16 synNotch receptor. CII-synNotch cells activated over 40x when plated on surfaces prepared with 25 μg/ml CII, whereas control LaG16-synNotch cells did not exhibit activation (**Figure 1E**). These results confirm that CII-synNotch enables specific recognition of CII substrates and regulates potent, inducible transgene expression.

### CII-synNotch MSCs accurately profile cartilage matrix degradation

Next, we transitioned from artificial, CII-decorated surfaces to more physiologically relevant substrates to test whether CII-synNotch enables cells to distinguish between healthy and damaged cartilage. First, we engineered a CII-synNotch MSC line to express the reporter transgene secreted alkaline phosphatase (SEAP) in response to ligand activation. Then, we harvested 3 mm-diameter cartilage biopsy punches from porcine limbs. We have previously shown that trypsin treatment of cartilage samples exposes collagen epitopes detected by mAbCII, mimicking the degradation of the ECM associated with arthropathy (33). Thus, we enzymatically treated a subset of the porcine explants with TrypLE Express for one hour to expose CII epitopes (**Figure 2A**). These and control explants were then washed in serum-containing PBS prior to seeding CII-synNotch MSCs on each explant. MSCs seeded on enzyme-treated explants produced higher levels of synNotch-driven SEAP compared to cells cultured on untreated explants (**Figure 2B**), suggesting that CII-synNotch cells are capable of profiling healthy versus degraded cartilage surfaces.

**Figure 2:**
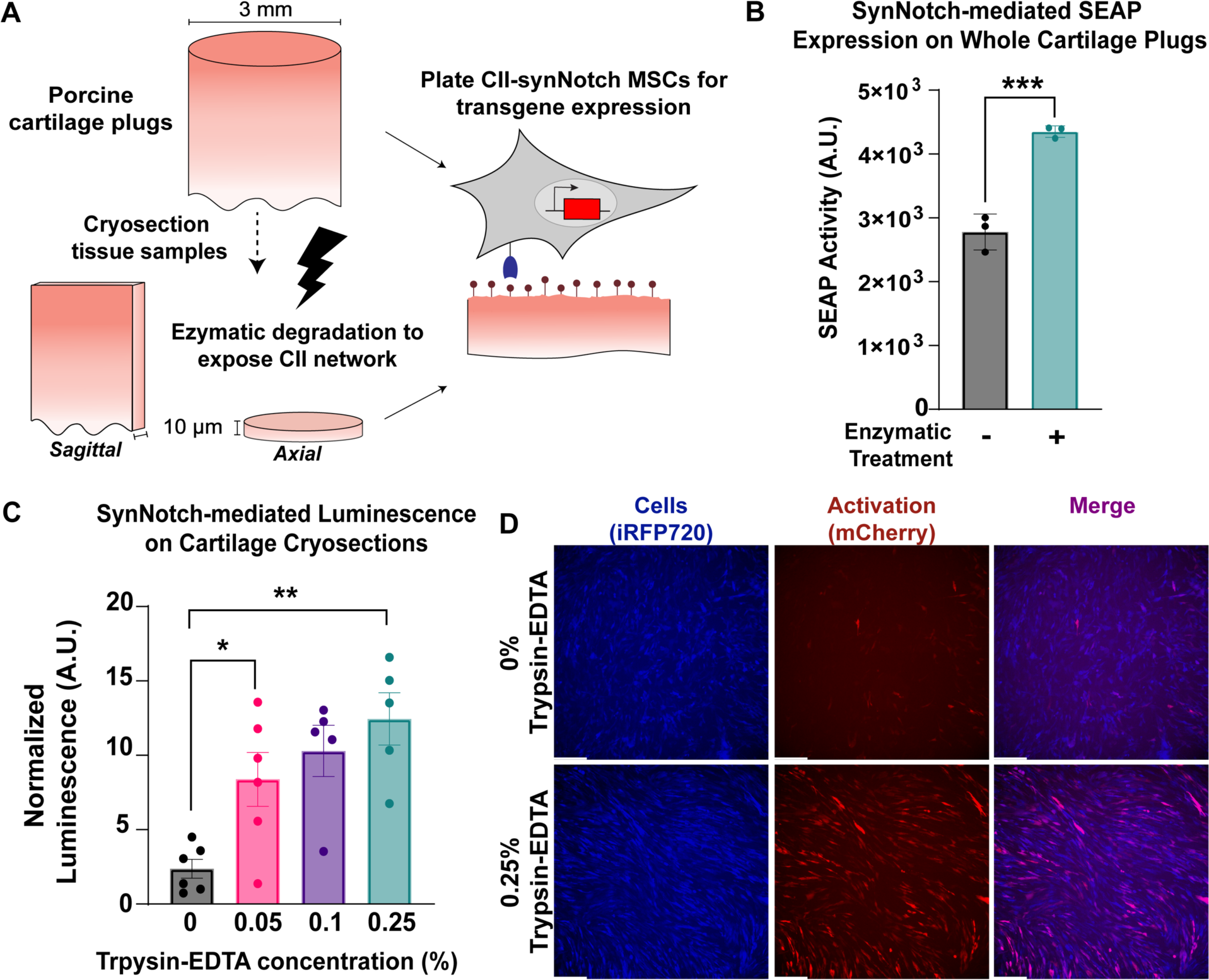
CII-synNotch MSCs detect degradation of cartilage tissue. **(A)** 3 mm-diameter primary porcine cartilage explants were enzymatically treated or cryosectioned prior to enzymatic treatment. Enzyme treatment reveals CII epitopes detected by the mAbCII recognition motif. **(B)** SynNotch-inducible SEAP expression after plating engineered cells on enzymatically-damaged whole-plug explants (t=72 hr, Student’s t-test on n=3 replicates per group). **(C)** Inducible luciferase expression from engineered cells on cryosections treated with varying concentrations of trypsin-EDTA (t=48hr). **(D)** Representative images of synNotch activation on sagittal cryosection samples without treatment and with 0.25% trypsin-EDTA (t=48hr). iRFP is false-colored as blue to enable reporter discrimination in the merged image. Scale = 200 µm. Data plotted as mean ± SEM. *p<0.05, **p<0.01.

While encouraging, we found it difficult to reliably seed MSCs on whole cylindrical explants. Cryosectioning the tissue specimen creates a consistently level surface for facile, reproducible cell seeding. Thus, we cryosectioned another subset of the explanted porcine cartilage into 10 μm thick sections both perpendicular to (axial) and parallel to (sagittal) the axis of the biopsied explant (**Figure 2A**). Axial sections were then treated with varying levels of trypsin-EDTA ranging from 0.05% to 0.25% prior to washes with serum-containing PBS and subsequent seeding of synNotch-MSCs capable of inducible luciferase expression. We found that trypsin treatment rendered a dramatic boost of synNotch-driven luciferase transgene expression (**Figure 2C**), even on samples treated with a low concentration of 0.05% trypsin-EDTA. Fluorescence microscopy reveals intense and uniform mCherry expression across a trypsin-treated sagittal section, whereas few cells are mCherry positive on an untreated section (**Figure 2D**). Some degree of matrix damage, and therefore mCherry signal, on such untreated sections is expected, due to the disruptive nature of collecting biopsy samples, cryosectioning them, and subjecting them to freeze-thaw cycles. Collectively, these results suggest that, like mAbCII itself (33, 35, 42), CII-synNotch MSCs accurately profile degraded versus control cartilage tissues.

### Engineered MSCs spatially constrain transgene expression to regions of CII exposure

Several synthetic biology approaches have been developed to gate expression of transgenes on the basis of inputs relevant to arthropathy, such as pro-inflammatory cytokines (25, 43) prominent in arthritic joints and mechanical load (26). The juxtacrine-like nature of synNotch requires physical immobilization of its ligand as a condition for activation. Thus, one potential advantage of CII-synNotch is the ability to spatially restrict transgene expression to regions defined by exposure of CII in the extracellular matrix. To determine whether this degree of spatial regulation may be lost by cells responding to type II collagen leached from arthritic cartilage (but not in direct contact with the tissue), we cultured CII-synNotch MSCs in control, untreated conditions, in the presence of 25 μg/ml CII solubilized in culture medium, or on CII-adsorbed surfaces (pre-treated with 25 μg/ml CII solution, as in prior experiments). Based on luminescence measures, there was only modest activation of luciferase expression when cells were cultured in the presence of 25 μg/ml CII solubilized in culture medium as compared to cells plated on adsorbed CII (**Figure 3A**). To mirror more representative conditions of leached collagen, we utilized our cryosection experimental configuration and monitored synNotch activation via fluorescence microscopy. Engineered MSCs were plated on trypsin-treated cryosections and allowed to proliferate and migrate beyond the margins of the cryosectioned specimen and throughout the culture well (**Figure 3B**). Fluorescence microscopy reveals that inducible mCherry transgene expression is abundant in cells in contact with the cartilage cryosection, whereas cells out of contact with the tissue remain mCherry-low (**Figure 3B**). Thus, CII-synNotch enables spatially regulated transgene expression induced by cellular contact with damaged ECM.

**Figure 3:**
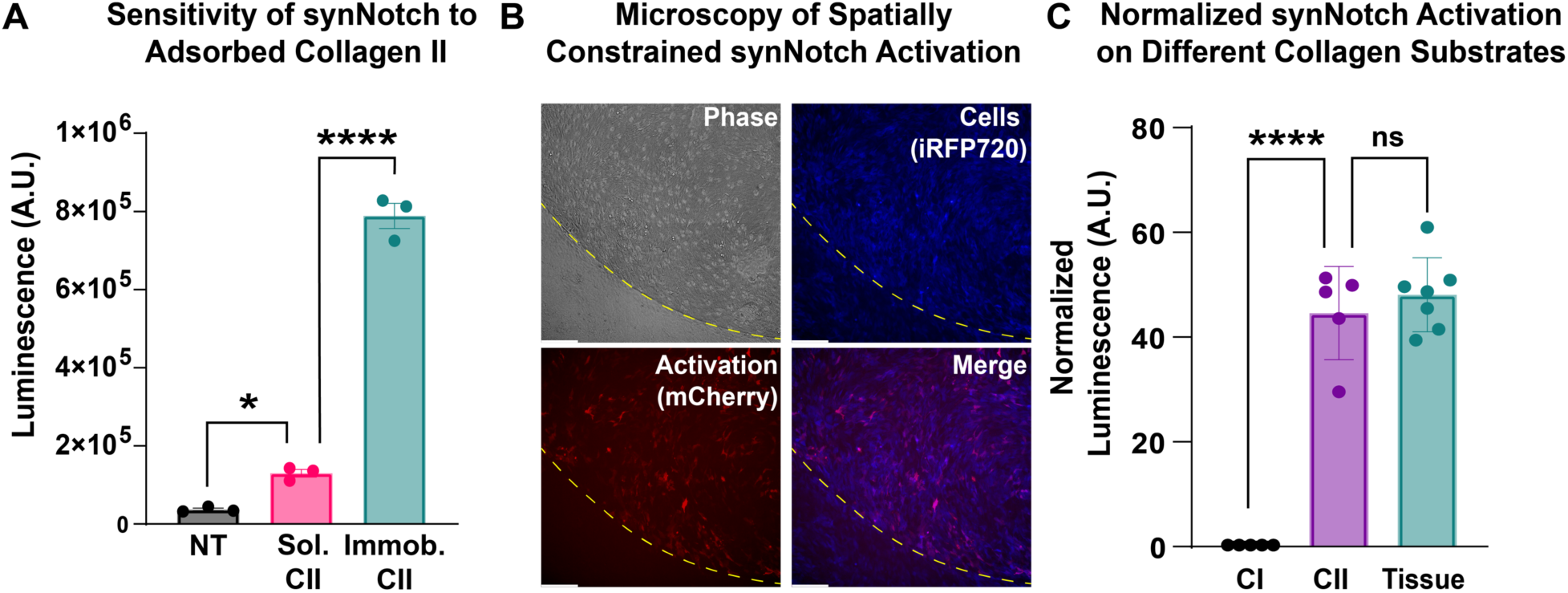
Transgene expression by CII-synNotch cells is spatially gated by direct contact with immobilized CII. **(A)** Luminescence activity from engineered MSCs cultured in untreated conditions, in medium supplemented with CII (“Sol. CII”) or on cell culture surfaces pre-treated with CII (“Immob. CII”) (t=72 hr). **(B)** Representative image of cartilage explant (outlined in yellow dash) containing CII-synNotch cells (blue) that express mCherry (red) upon activation. Scale = 200 µm. iRFP is false-colored as blue to enable reporter discrimination in the merged image. **(C)** Comparison of CII-treated surfaces and trypsinized cryosectioned cartilage tissue, as measured with normalized luciferase (t=48hr). Scale = 200 µm. Data: n≥3 replicates, plotted as mean ± SEM. Statistical analysis: one-way ANOVA with Tukey’s post-hoc analysis, *p<005, ****p < 0.0001. NT: no treatment

Finally, in preparation for experiments designed to assess the regenerative potential of CII-synNotch cells, we sought to validate the use of CII-decorated surfaces as a surrogate for damaged cartilage as an activating input for synNotch MSCs. Thus, we prepared culture surfaces treated with 25 μg/ml CI or CII as well as trypsin-treated cryosections. We then plated CII-synNotch MSCs on these culture substrates. After three days, we assessed synNotch-regulated firefly luciferase activity. Consistent with prior results, we again saw a drastic, 48-fold increase in synNotch signaling when cells were cultured on CII versus CI (**Figure 3C**). Additionally, we saw comparable levels of transgene expression between cells cultured on 25 μg/ml CII-treated surfaces versus enzymatically damaged cartilage. These results indicate that surfaces treated with 25 μg/ml CII closely approximate the epitope density that CII-synNotch cells detect in degraded cartilage tissue. These data support the use of CII-decorated substrates in experiments that evaluate CII-synNotch cell performance in cartilage regenerative medicine.

### CII-synNotch MSCs engineered for pro-anabolic transgene expression

MSCs are of therapeutic interest in arthritis because, in addition to their differentiation potential (44–46), they are strong paracrine signalers that can secrete trophic factors aiding in tissue preservation (47, 48). One such growth factor is transforming growth factor-β3 (TGFβ3), which promotes chondrocyte proliferation, reduces chondrocyte apoptosis, and enhances expression of matrix constituents critical for cartilage function, such as CII and aggrecan (49, 50). We engineered a CII-synNotch MSC line expressing a synNotch payload cassette encoding human *TGFβ3* coupled with reporter mCherry expression via an internal ribosome entry site (IRES). In validating the performance of this cell line, flow cytometry results revealed dramatic, CII-dependent upregulation of mCherry in synNotch cells cultured on CII-decorated surfaces (**Figure 4A**). Furthermore, ELISA measurements of inducible TGFβ3 protein production reveal significantly augmented levels of the growth factor in the presence of immobilized CII, with secreted TGFβ3 levels being near or below the detection limit in the control conditions but approximately 6 ng/ml in synNotch-active conditions (**Figure 4B**). As TGFβ3 production is closely linked to improvements in cartilage ECM composition (50, 51), qRT-PCR was performed to profile genes associated with matrix synthesis. As expected, transgene human *TGFβ3* levels were inducibly expressed in the synNotch-TGFβ3+mCherry cell line but not in a synNotch-luciferase+mCherry control (CTRL) reporter cell line (**Figure 4C**). Notably, mouse *Acan* and *Col2a1* genes were upregulated in response to CII-induced, synNotch-mediated TGFβ3 expression, whereas CTRL cells did not demonstrate the same levels of *Acan* and *Col2a1* upregulation (**Figure 4D**). *Col1a1* expression was also reduced in a CII and cell-line-dependent manner. These results demonstrate the capacity of synNotch-engineered cells to govern the expression of cartilage anabolic factors that positively influence signatures of matrix synthesis in an exposed CII-dependent manner.

**Figure 4:**
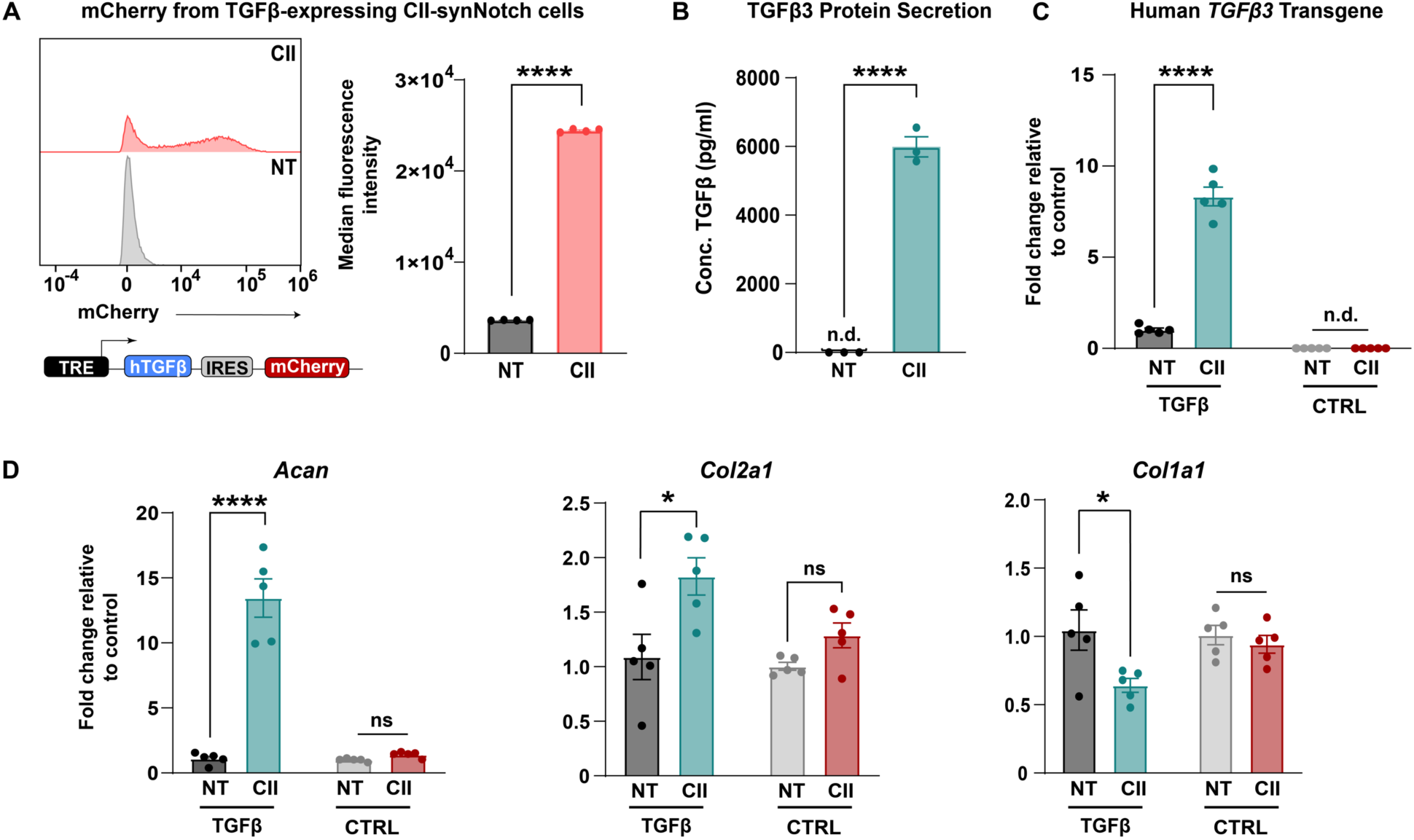
CII-synNotch effectively regulates pro-anabolic TGFβ3 expression. **(A)** A synNotch payload contains human TGFβ3 co-expressed with mCherry, connected through an internal ribosomal entry site (IRES), allowing for expression of both transgenes. Inducible mCherry measurements of TGFβ3-expressing cells with and without activation on CII as a readout of synNotch activation (t=72 hours, Student’s t-test on n=3 replicates per group). **(B)** ELISA measurements of TGFβ3 after plating on CII-adsorption to cell culture surfaces (t=72 hours, Student’s t-test on n=3 replicates per group). **(C)** qRT-PCR detection of transgene *hTGFβ3* in experiments comparing TGFβ3-expressing cells to reporter-expressing cells. qRT-PCR data is normalized to no-treatment controls for each cell line. **(D)** qRT-PCR detection of pro-anabolic *Acan* and *Col2a1* expression, as well as *Col1a1* expression. All qRT-PCR: t=72 hrs, two-way ANOVA with Tukey’s post hoc analysis on n=5 replicates per group. *p<0.05, ****p<0.0001. n.d.: not detected.

### CII-synNotch MSCs attenuate inflammation experienced by chondrocytes

There is an established correlation between dysregulated inflammatory cascades (52–54) and articular cartilage catabolism. As such, mitigating activity of pro-inflammatory cytokines would aid in the amelioration of arthritic lesions. We have previously demonstrated the ability of GFP-sensitive synNotch MSCs to programmably secrete the transgene soluble tumor necrosis factor receptor type 1 (sTNFR1), which mitigates inflammation induced by TNF-α (10, 31). Another OA-associated pro-inflammatory cytokine implicated in tissue breakdown is IL-1, which is mediated in part by activation of the master transcription factor nuclear factor kappa B (NF-κB) (55). With the goal of generating a synthetically regulated MSC line capable of mitigating an inflammatory environment in response to cartilage damage, CII-synNotch MSCs were engineered with inducible human IL-1 receptor antagonist (IL-1Ra). IL-1Ra is an established anti-inflammatory biologic that is clinically used in the treatment of rheumatoid arthritis (56), and has demonstrated preclinical benefits in the treatment of OA (25, 43, 57–60). Thus, we engineered a synNotch-inducible IL-1Ra MSC line. As was the case with the TGFβ3 cell line, synNotch-regulated IL-1Ra expression was dramatically induced when cells were cultured on CII-decorated surfaces (**Figure 5A**).

**Figure 5:**
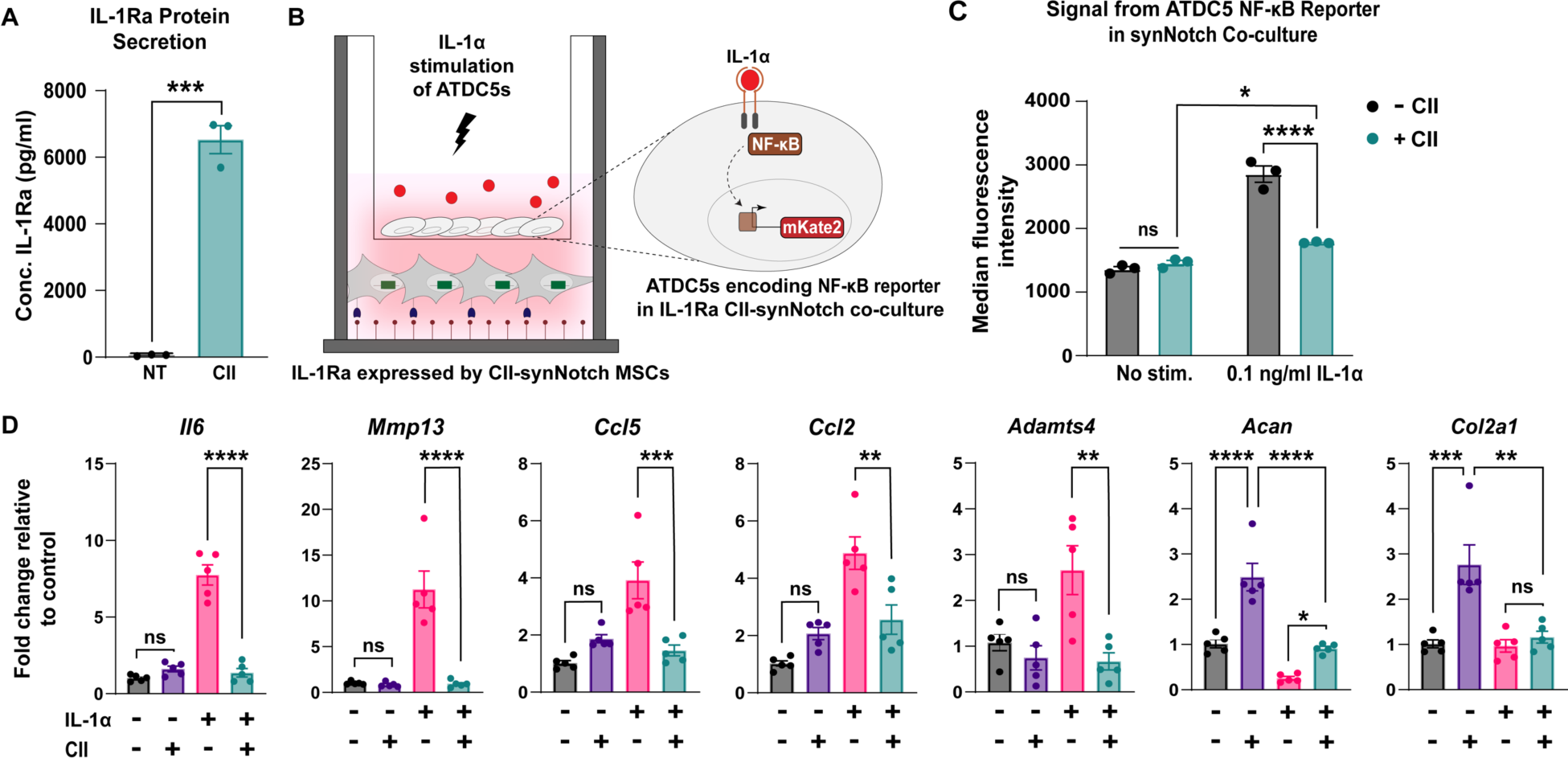
CII-synNotch MSCs mitigate IL-1-induced inflammation in a CII-dependent manner. **(A)** ELISA measurements of IL-1Ra after plating on CII-decorated surfaces (t=72 hours, Student’s t-test on n=3 replicates per group). **(B)** Schematic of trans-well co-culture for assessment of IL-1Ra synNotch circuits on inflammatory chondrocytes. Bottom: 25 µg/ml of CII is plated for activation of CII-synNotch MSCs containing inducible IL-1Ra. Top: ATDC5 chondrocytes are plated in 3 µm-porous inserts. After three days of synNotch activation by MSCs, 0.1 ng/ml IL-1α was added to the insert. Flow cytometry and qRT-PCR results were obtained two days after stimulation. **(C)** ATDC5s were transduced with an mKate2 reporter that is upregulated in response to NF-κB-driven inflammation. Flow cytometry of mKate2 reports for the effect of inducible IL-1Ra expression on the NF-κB signal in IL-1-stimulated co-cultures (t=5d from plating, two-way ANOVA on n=3 replicates per group). **(D)** qRT-PCR detection of pro-inflammatory genes *Il6, Mmp13, Ccl5, Ccl2,* and *Adamts4* as well as ECM-related *Acan* and *Col2a1.* All qRT-PCR data is normalized to chondrocyte gene expression in co-cultures with neither CII-synNotch activation nor IL-1α stimulation (t=5d from plating, two-way ANOVA on n=five replicates per group). *p<0.05, **p<0.01, ***p<0.001, ****p<0.0001.

These synNotch cells were then tested in a trans-well co-culture with mouse ATDC5 chondrocytes containing a fluorescent mKate2 reporter of NF-κB transcriptional activity (37) (**Figure 5B**). SynNotch-IL1Ra MSCs were plated on CII- or untreated surfaces in the bottom chambers of trans-wells, and chondrocytes were plated in upper chambers on inserts with 3 μm pores. Three days later, reporter chondrocytes were stimulated with 0.1 ng/ml IL-1α. Flow cytometry of NF-κB reporter chondrocytes reveal a synNotch-regulated reduction in mKate2 levels (**Figure 5C**), reflecting reduced inflammatory transcriptional activity in the IL-1-treated chondrocytes co-cultured with CII-responding, synNotch-IL1Ra MSCs. To determine the influence of synNotch-regulated IL-1Ra on chondrocyte gene expression, we performed qRT-PCR on mRNA isolated from chondrocytes in the upper chamber of trans-wells. Notably, several genes implicated in the breakdown of articular cartilage, such as the cytokine *Il6*, matrix-degrading enzymes *Mmp13* and *Adamts4*, and chemokine ligands *Ccl5* and *Ccl2*, were upregulated in chondrocytes upon IL-1α stimulation when synNotch-MSCs were cultured in the CII-free “off” state (**Figure 5D**). Additionally, matrix synthesis genes such as *Acan* and *Col2a1* were markedly reduced in chondrocytes by IL-1α treatment, as expected (61–63). However, when synNotch-MSCs were activated by CII to produce IL-1Ra, the inflammatory transcripts *Il6*, *Mmp13*, *Ccl5*, *Ccl2* and *Adamts4* were significantly attenuated. Furthermore, there was partial restoration of *Acan* when cells were cultured in the CII-including “on” state. Given the upregulation in *Acan* in the IL-1(-)/CII(+) condition, part of the *Acan* restoration may be attributable to the influence of CII on *Acan* expression rather than synNotch-driven IL1Ra production. Taken together, these findings indicate that synNotch-mediated IL-1Ra expression protects chondrocytes from the deleterious impact of supraphysiologic IL-1 treatment and may serve to stabilize the joint in the face of inflammation that drives arthritis progression.

## DISCUSSION

A lack of clinically available DMOADs motivates the need for improving efficacy of prospective arthritis cell-based therapies. With only few exceptions, evidence does not indicate that minimally manipulated cell-based therapies intrinsically sense and resolve arthropathy. Deployment of cells to mitigate inflammation and tissue catabolism implicated in OA has presented challenges such as unregulated expression of pleiotropic factors used as OA therapeutics as well as cartilage hypertrophy (64, 65). We and others have developed approaches to instruct cell behaviors to execute reliable cartilage regenerative functions in an effort to overcome limitations of minimally manipulated cells or to circumvent drawbacks of traditional genetic engineering (11, 26, 63, 66). Prior approaches rewire native signaling pathways to enable cells to autonomously regulate therapeutic drug expression. For example, we previously inserted transgenes such as sTNFR1 and IL-1Ra to the *Ccl2* locus of stem cells to govern inflammation-inducible expression of these anti-inflammatory factors in stem cell-derived chondrocytes (25, 67). However, such rerouted output from the *Ccl2* locus is susceptible to crosstalk, as several inputs independent of arthropathy activate *Ccl2* expression. Further, these routes of signaling do not guide cells to execute regenerative functions restricted to regions of cartilage damage. This means the feedback-controlled expression offered by such design strategies is neither uniquely localized to the joint nor specific to OA.

By contrast, here we demonstrate the power of an artificial receptor with engineered specificity for cartilage degradation that operates in a manner that is orthogonal to native cell signaling pathways. The synNotch platform was previously established to enable artificial signaling to regulate epithelial patterning (27), to improve CAR T cell discriminatory power and function (28, 68), and to specify human pluripotent stem cell differentiation (31). The modular nature of synNotch facilitates the design of a synthetic receptor that enables a cell to recognize damaged joint structures to subsequently implement defined, predetermined behaviors. By exploiting an scFv derived from mAbCII as a recognition motif for synNotch, we licensed MSCs to uniquely identify OA-associated cartilage degradation, constraining therapeutic drug production to instances where CII fibers are exposed in damaged ECM, as opposed to nonspecific conditions such as generalized inflammation. This feature can serve to minimize unnecessary transgene expression that can disrupt normal tissue homeostasis. Thus, the CII-synNotch platform is distinct from existing approaches, because it enables customized transgene expression that is spatially constrained to sites of cartilage degradation.

A major goal of our study was to determine the specificity and dynamic range of CII-synNotch signaling. Engineered cells demonstrated robust activation on immobilized CII, with peak activation attained when surfaces were prepared with a CII solution of 25 μg/ml. Crucially, CII-synNotch MSCs did not activate on surfaces decorated with collagen type I, suggesting specificity of the receptor that mimics that of mAbCII itself. Indeed, even in conditions where cells were exposed to the same concentration of CII, but in solution as opposed to surface-adsorbed collagen, artificial CII-synNotch signaling was only modestly induced, reflecting the juxtacrine-like behavior of synNotch that requires ligand immobilization. Importantly, we found that CII-synNotch activation from a solution prepared by passive adsorption of a 25 μg/ml CII corresponded closely to that attained via the CII network presented by degraded cartilage cryosections. Validation of such CII-decorated surfaces as surrogates for tissue specimen simplified subsequent studies of cell behaviors regulated by CII-synNotch, which showed exquisite regulation of TGFβ3 and IL-1Ra, demonstrating a wide dynamic range spanning undetected levels in synNotch-off conditions to levels of ng/ml produced by activated synNotch-MSCs.

This range of inducible growth factor expression potentiates customized engineering of MSC behaviors for cartilage regenerative medicine. Efforts to stimulate ECM synthesis and reverse OA pathology via TGFβ overexpression are under investigation in human clinical trials, as with the Kolon TissueGene “TG-C” product (69). However, high levels of this growth factor may also result in suppressed immune cell function, cartilage hypertrophy, and endochondral bone formation (64, 70, 71). Using the CII-synNotch receptor to drive inducible TGFβ expression may mitigate negative side effects of excessive TGFβ by creating a sense/response pipeline that governs tunable growth factor expression. Our results illustrated the capacity to produce significant levels of TGFβ3 protein by MSCs that positively influenced expression of genes associated with ECM synthesis. In particular, synNotch-MSCs produced ∼6 ng/ml of TGFβ3, on par with prior reports of ectopic, constitutive TGFβ3 production after lentiviral transduction of MSCs (51) and also approximating the 10 ng/ml levels commonly used for *in vitro* cartilage tissue engineering. Here, these levels were adequate to enhance expression of *Acan* and *Col2a1*, which encode for major cartilage ECM constituents. Additionally, the reduction of *Col1a1* in a synNotch-dependent manner further supports the development of a pro-chondrogenic phenotype of these cells, despite the multipotent capacity of MSCs to differentiate into cells of CI-rich subchondral bone (44). Future work with anabolic synNotch circuits will assess the effect of selective TGFβ3 on ECM production in the context of three-dimensional engineered tissues and will also explore alternative matrix-stimulating, synNotch-regulated transgenes, including GDF5 as well as members of the FGF and IGF families.

In addition to demonstrating the capacity of synNotch-MSCs to regulate cartilage anabolic activities, we also evaluated whether CII-synNotch cells could selectively antagonize inflammation that is featured in OA. IL-1 is a prominent player in OA pathogenesis, particularly in post-traumatic OA (PTOA), and increased levels in serum and synovial fluid correlate with OA incidence and severity (58, 72, 73). Viral delivery of IL-1Ra has improved outcomes of PTOA in large animal models, but safety concerns for delivery to the joint include increased infections due to prolonged IL-1 suppression and perivascular cuffing secondary to viral gene therapy (57, 74, 75). Here, IL-1Ra-expressing CII-synNotch cells modulated the IL-1α-induced inflammatory response of ATDC5 chondrocytes, as measured by NF-κB transcriptional activity and qRT-PCR profiling of several inflammation-responsive genes. We also detected a trend in restoring ECM-associated gene expression for *Acan* after both synNotch activation and IL-1 stimulation. However, the presence of CII in the co-culture positively influenced *Acan* expression, suggesting further investigation is required to attribute *Acan* restoration to synNotch-regulated IL-1Ra expression. Future extension of this work will establish the ability of the CII-synNotch platform to antagonize other degradative factors in the joint, including matrix-degrading enzymes and pro-inflammatory activity of the cytokines IL-6, IL-17, and TNF.

A synthetic, arthritis-sensitive receptor presents a next-generation approach for the treatment of arthropathies via engineered cell-based therapies. Other molecular therapies, such as siRNA or *in situ* delivery of CRISPR-based reagents (76), offer alternative strategies to modulate OA gene regulation and can be combined with targeting moieties such as mAbCII (35, 77–79). While siRNA and CRISPR reagents can be targeted to any gene, they are not responsive to arbitrarily selected inputs, such as damaged ECM components, that cells can be engineered to recognize. By programming cells with synNotch, they are endowed with the ability to sense pre-selected inputs to execute user-specified functions without the need for introducing activating reagents, as is the requirement for drug-inducible gene therapies. Furthermore, CII-synNotch cells integrate the benefits of gene therapy with synthetic biology to create designer cells as candidate disease-modifying agents that can carry out diverse assignments including matrix synthesis, amelioration of inflammation, and recruitment of progenitors via chemokine production.

When considering the potency of CII-synNotch cells *in vivo*, transgenes need to restore cartilage integrity and resolve joint inflammation. Our workflow for generation of multiple candidate payloads provides a platform for facile design of combinatorial transgene delivery that may mitigate destructive factors that define the arthritic microenvironment. For example, creating a multicistronic CII-synNotch payload cassette with both pro-anabolic and anti-catabolic factors (e.g., co-expression of TGFβ3 and IL-1Ra) could improve efficacy of engineered MSCs. Future studies will test such designer cells *in vitro* and in mouse models of spontaneous OA and PTOA, such as the STR/ort mouse model (80, 81) and the anterior cruciate ligament rupture model (82, 83), respectively. Further, the destructive nature of rheumatoid arthritis (RA), and the ability of mAbCII to identify such RA-affected tissue (34, 84), implies that CII-synNotch cells may potentiate therapeutic outcomes in the context of RA, as well.

In summary, we used synthetic biology to program primary cells with an artificial receptor containing an OA-specific recognition motif. In an integrated platform, we leveraged the specificity of the mAbCII antibody and exploited the versatility and juxtacrine-like nature of synNotch to engineer MSCs for disease-dependent, autoregulated expression of transgenes that may mitigate arthritis. This work lays the foundation for continued investigation of synNotch to develop cell-based DMOADs.

## Acknowledgements

This work was supported by the Arthritis National Research Foundation Judy E. Green Valiant Women’s Fellowship (JMB), National Institute of Arthritis and Musculoskeletal and Skin Diseases 1R21AR079683 (JMB, CLD, KAH), National Science Foundation Faculty Early Career Development Program (CAREER) CBET-2237639 (JMB), VA Merit Award BX005195 (KAH), and National Science Foundation Graduate Research Fellowship (BLW). Cell sorting was performed in the Vanderbilt Flow Cytometry Shared Resource, which is supported by the Vanderbilt Ingram Cancer Center (P30CA068485) and the Vanderbilt Digestive Disease Research Center (DK058404). We also acknowledge the Vanderbilt University Translational Pathology Shared Resource core, supported by NCI/NIH Cancer Center Support Grant P30CA068485, for all tissue sectioning and processing. The authors thank Mr. Saurav Bhattarai, Mr. Zachary Eidman and Mr. Jiageng Liu for technical assistance.

